# Global diversity and dispersal routes of the *Ostreid herpesvirus type 1* infecting *Magallana gigas*

**DOI:** 10.64898/2026.06.05.730385

**Authors:** Camille Pelletier, Germain Chevignon, Maude Jacquot, Benjamin Morga

## Abstract

The order Herpesvirales comprises double-stranded DNA viruses characterized by substantial genomic plasticity, including recombination, structural variation, gene gain and loss, and lineage turnover. These processes can obscure phylogenetic relationships and complicate the reconstruction of viral evolutionary histories. Within this order, Ostreid herpesvirus 1 (OsHV-1) is a major pathogen of the Pacific oyster *Magallana gigas* and is responsible for recurrent mortality events affecting global aquaculture. Early molecular investigations based on partial genomic regions identified several viral lineages, including the “var” and “µVar” lineages, but provided limited resolution for genome-wide evolutionary inference. The subsequent availability of complete genomes revealed extensive structural variation, such as insertions, deletions, and genomic rearrangements, highlighting the high genomic plasticity of OsHV-1. Although phylogenomic analyses have estimated evolutionary rates compatible with other large double-stranded DNA viruses, current inferences remain based on geographically restricted datasets, leaving the global evolutionary dynamics of OsHV-1 within its principal host insufficiently resolved. Here, we present 275 newly sequenced OsHV-1 genomes collected from infected *M. gigas* oysters between 1994 and 2022 across major oyster-producing regions worldwide. Using *de novo* genome assembly combined with comparative genomics, population genetic analyses, and time-scaled phylogenetic reconstruction, we investigate global genomic diversity and the spatio-temporal dynamics of viral diversification. Our results reveal long-standing viral diversity in East Asia, the emergence of structurally distinct Pacific and microvariants lineages, and ongoing diversification shaped by recombination, structural genome plasticity, and anthropogenic oyster movements. By integrating three decades of whole-genome data, this study provides a phylogenomic framework for understanding the diversity, evolution, and dispersal of OsHV-1 in modern aquaculture systems.

## Introduction

Herpesvirales is a diverse order of large double-stranded DNA viruses infecting vertebrate and invertebrate hosts. Beyond their medical and veterinary relevance, herpesviruses are increasingly recognized as dynamic evolutionary systems characterized by recombination, structural genome variation, gene gain and loss, and lineage turnover (Brown & Smith, 2018; Dellicour et al., 2023; Dotto-Maurel et al., 2025; Houldcroft et al., 2017). Whole-genome sequencing has shown that herpesvirus diversification often involves recurrent insertions and deletions, genomic rearrangements, and mosaic ancestry resulting from recombination (Norberg et al., 2007; Ortigas-Vasquez & Szpara, 2024; Pérez-Losada et al., 2015; Szpara et al., 2014). These processes can obscure simple phylogenetic relationships and complicate inference of transmission history, highlighting the importance of high-resolution phylogenomics to reconstruct evolutionary trajectories (Nater et al., 2015; Rannala, 2025).

Within the family *Malacoherpesviridae*, Ostreid herpesvirus type 1 (OsHV-1; species *Ostreavirus ostreidmalaco1*) is one of the most consequential viral pathogens affecting marine invertebrates (Arzul et al., 2017; Davison et al., 2005; Farley et al., 1972; Nicolas et al., 1992; Segarra et al., 2010). OsHV-1 is associated with recurrent mass mortality events in the Pacific oyster *Magallana gigas*, which represents nearly 98% of global oyster production worldwide (FAO, 2023). Since its initial detection in the early 1970s (Farley et al., 1972; Nicolas et al., 1992), the virus has been reported across Europe, Asia, Oceania and North Africa (Batista et al., 2015; Hwang et al., 2013; Jenkins et al., 2013; Lynch et al., 2012; Moss et al., 2007; Peeler et al., 2012). Its global dissemination likely parallels the historical expansion of oyster aquaculture. Indeed, large-scale international transfers of spat and broodstock during the twentieth century, combined with repeated movements among production areas within farming cycles, have created extensive host connectivity and potential pathways for viral dispersal (Grizel & Heral, 1991; Lupo et al., 2016; Nell, 2001; Wörner et al., 2019). These anthropogenic processes are expected to have profoundly influenced the global phylogeography of OsHV-1.

Despite its detection in several bivalve species, including scallops and clams (Arzul, Nicolas, et al., 2001; Arzul, Renault, et al., 2001; Bai et al., 2019; Renault & Arzul, 2001; Xia et al., 2015), experimental evidence demonstrating that OsHV-1 can complete its infectious cycle in species other than *M. gigas* is still lacking. *M. gigas* remains the dominant host species in terms of global production and disease impact. Consequently, understanding viral diversification within *M. gigas* populations is central to reconstructing OsHV-1 evolutionary history and epidemiology.

Early molecular investigations based on PCR amplification and targeted sequencing of short genomic regions identified distinct viral variants, including the “var” lineage and later the “µVar” group associated with massive mortality outbreaks in Europe from 2008 onward (Arzul, Renault, et al., 2001; Martenot et al., 2011; Segarra et al., 2010). However, these variants were defined using a limited number of loci and provided restricted resolution for inferring genome-wide relationships. The publication of the OsHV-1 reference genome in 2005 (Davison et al., 2005) and subsequent complete genomes of microvariants (Burioli et al., 2017) enabled comparative genomic analyses at a broader scale. These studies revealed recurrent structural variation, including large deletions and insertions, particularly within the unique long (U_L_) region (Abbadi et al., 2018; Bai et al., 2019; Burioli et al., 2017; Delmotte-Pelletier et al., 2022; Xia et al., 2015). Such structural polymorphisms can affect coding regions and gene content, suggesting potential functional consequences and highlighting genome plasticity as a key feature of OsHV-1 evolution.

Recombination represents another important evolutionary mechanism shaping herpesvirus diversity. Phylogenomic analyses in several herpesvirus systems have demonstrated that recombination can generate mosaic genomes and accelerate lineage diversification (Dellicour et al., 2023; Houldcroft et al., 2017). In OsHV-1, signatures of recombination have been detected in genome-wide analyses, although their frequency, distribution and epidemiological relevance remain incompletely quantified. Structural variation and recombination together may contribute to rapid diversification, lineage emergence and adaptation within host populations (LaTourrette & Garcia-Ruiz, 2022; Montoya et al., 2021).

Recent phylogenomic work has also begun to address the temporal dimension of OsHV-1 evolution. Estimates of substitution rates based on time-calibrated genomic datasets indicate that OsHV-1 evolves at rates consistent with other large double-stranded DNA viruses, while still allowing measurable evolution over epidemiological timescales (Delmotte-Pelletier et al., 2022; Morga-Jacquot et al., 2021). These findings support the feasibility of reconstructing recent divergence events and dispersal histories. However, rate estimates remain based on limited sample sizes and geographically restricted datasets, leaving uncertainty regarding the robustness of temporal signal and the generality of inferred evolutionary dynamics at the global scale.

Recent phylogenomic evidence further suggests that host-associated processes can influence OsHV-1 diversification. Pelletier et al. (2025) reported genetic divergence between viral populations infecting *M. gigas* and *Ostrea edulis*, supporting the possibility of host specialization. These findings emphasize that ecological and host-related factors may shape viral population structure. However, the global evolutionary dynamics of OsHV-1 within its principal aquacultured host, *M. gigas*, remain insufficiently resolved.

Despite the growing number of publicly available genomes (approximately one hundred complete assemblies), sampling remains geographically and temporally uneven (Bai et al., 2019; Battistel et al., 2025; Davison et al., 2005; Delmotte-Pelletier et al., 2022; Pelletier et al., 2025; Xia et al., 2015). Only two studies have explicitly examined spatio-temporal genomic patterns. One relied on read-mapping approaches that may bias the detection of insertions and low-frequency variants (Morga-Jacquot et al., 2021), while the other, based on *de novo* assemblies, identified regional clustering and suggested transmission linked to oyster movements but was limited in sample size (Delmotte-Pelletier et al., 2022). As a result, fundamental questions remain unresolved: To what extent are OsHV-1 populations infecting *M. gigas* structured at the global scale? Does contemporary diversity reflect a limited number of ancestral introductions followed by regional diversification? What is the relative contribution of recombination and structural genome variation to lineage emergence? And how have anthropogenic oyster transfers influenced viral phylogeography over the past three decades?

Addressing these questions requires a globally representative dataset combined with high-resolution phylogenomic analyses. Here, we present 275 newly sequenced OsHV-1 genomes sampled from *M. gigas* between 1994 and 2022 across major oyster-producing regions worldwide. Using *de novo* assembly and integrated comparative genomics, population genetic and phylogenetic approaches, we aim to *(i)* characterize global genomic diversity within the dominant host species, *(ii)* reconstruct evolutionary relationships and divergence patterns among lineages, and *(iii)* investigate the spatio-temporal dynamics of viral diversification in the context of worldwide oyster trade. By focusing on the principal aquaculture host, this study provides a comprehensive framework to understand the evolutionary processes shaping OsHV-1 diversity and persistence at the global scale.

## Results

A total of 275 OsHV-1–infected *M. gigas* samples, collected worldwide between 1993 and 2022, primarily from France (n=238) and eight from other countries (Ireland, Korea, New Zealand, Germany, Australia, Spain, USA), were sequenced and subjected to *de novo* assembly using a standardized bioinformatic pipeline. The reconstructed genomes were analyzed for structural variations, while SNPs were used to infer population structure and visualize admixture patterns. Genetic relationships among lineages and patterns of viral diversification across geographic regions and time were further explored using network analyses and phylogenetic approaches.

### Structural variations

A whole genomic structural analysis was performed using the alignment of the 275 *de novo* assembled Non-Redundant genomes (NR-genomes, used to simplify the OsHV-1 genome structure by retaining a single copy of each repeated region in the viral sequence, Delmotte-Pelletier et al., 2022) from worldwide samples together with four publicly available OsHV-1 genomes (NC_005881, Davison et al., 2005; KY242785 and KY271630, Burioli et al., 2017; MG561751, Abbadi et al., 2018). This global comparison revealed six distinct Genomic Architectures (GA), defined by their shared structural features relative to the OsHV-1 reference genome (NC_005881.2; Davison et al., 2005) (**Erreur ! Source du renvoi introuvable.**, TableS3).

GA 1 (n = 9; France, 1993–2008) comprises genomes closely matching the reference architecture (NC_005881.2, Davison et al., 2005), with no large-scale rearrangements and an average NR-genome length of ∼190 kbp.

GA 2 (n = 11; France, 2005–2008) is characterized by a single 8.5-kbp inversion in the U_L_ region between Open Reading Frame (ORF) 36 and ORF 43 and by two small insertions at positions 61038 (U_L_) and 184204 (IRS), which represent the earliest detectable structural divergences from the reference genome. This region includes ORFs encoding a membrane domain-containing protein (ORF 36), a transmembrane protein (ORF 41), two zinc-finger, ring-type domain proteins (ORF 38 and ORF 42), a secreted protein (ORF39), and three proteins of unknown function (ORF 37, ORF 40, and ORF 43).

GA 3 (n = 232; Europe, 2008–2022) corresponds to the well-established µVar lineage (Burioli et al., 2017). These ∼204 kbp genomes exhibit ∼10 kbp of insertions and deletions resulting in the loss of eight ORFs (ORF 11, ORF 36, ORF 37, ORF 38, ORF 48, ORF 62, ORF 63, and ORF 114) and the gain of four insertion elements (ORF IN.1, ORF IN.2, ORF IN.3, ORF IN.4), thereby defining a structural configuration characteristic of the µVar lineage (Burioli et al., 2017; Segarra et al., 2010). The deleted ORFs encode proteins involved in membrane association, zinc-finger motifs, and unknown functions.

GA 4 (n = 3; New Zealand, 2011) retains the µVar genomic framework but harbors a ∼420 bp insertion within ORF 115 and four additional deletions: two ∼60 bp deletions in the US, a ∼450 bp deletion within ORF 117 at the junction between X and IRS and finally a ∼5.4 kbp deletion spanning the IR_S_-U_S_ junction between ORF 120 and ORF 123, which together distinguish this lineage from classical µVar genomes.

GA 5 (n = 15; Tomales Bay, New Zealand, Australia, 2006–2018) shows a distinct Pacific-associated architecture. These ∼184 kbp genomes contain a 4-kbp inverted translocation at the right boundary of the IR_S_ region (ORF 121 and ORF 122), repositioning the U_S_ region internally within the IR_S_. They also carry four insertions: a 2.6-kbp insertion already described for µVar lineages, a ∼420 bp insertion in the ORF 115 already described in GA4, a ∼90 bp and ∼460 bp within intergenic regions; and seven deletions ranging from 62 bp to 1.5 kbp and one of 446 bp across the U_L_ (ORF 36, ORF 37, ORF 38, ORF 48), the IR_s_ (ORF 117) and U_S_ (ORF 123) regions respectively. Moreover, they possess a 6-bp deletion in the microsatellite region in place of the µVar-specific 12-bp deletion. These genomes have been reduced from ∼13 kbp.

GA 6 (n = 5; San Diego, Korea, 2012–2018) displays a distinct Pacific-associated genomic architecture. These ∼184 kbp genomes contain a 4-kbp inverted translocation at the right boundary of the IR_S_ region (ORF 121 and ORF 122), which repositions the U_S_ region internally within the IR_S_ (**Erreur ! Source du renvoi introuvable.**). They also carry four insertions: a 2.6-kbp insertion previously described in µVar lineages, a ∼420 bp insertion within ORF 115 already reported in GA4, as well as two additional insertions of ∼90 bp and ∼460 bp located in intergenic regions. In addition, GA6 genomes harbor seven deletions ranging from 62 bp to 1.5 kbp, as well as a 446-bp deletion, affecting loci distributed across the U_L_ (ORF 36, ORF 37, ORF 38, ORF 48), IR_S_ (ORF 117), and U_S_ (ORF 123) regions. They also exhibit a 6-bp deletion in the microsatellite region, replacing the µVar-specific 12-bp deletion. Altogether, these structural rearrangements have resulted in a cumulative genome reduction of ∼13 kbp. All the ORFs affected by these rearrangements correspond to those previously described.

### Population structure, admixture and recombination of OsHV-1 genomes

The population admixture and genetic structure analysis, performed with STRUCTURE (Raj et al., 2013) and using multiple independent runs of the Gibbs sampler, indicated that the optimal number of ancestral populations in the dataset was K = 9. Each OsHV-1 genome was assigned to one or more of these ancestral populations, based on population admixture and similarity of ancestral populations, allowing us to define four arbitrary admixed populations (*i.e.* assignment to pre-defined populations) in a temporally and structurally coherent framework (**Erreur ! Source du renvoi introuvable.**).

The first admixed population is composed of older French genomes, collected before 2008, corresponding to the reference lineage, and genomes collected in the Pacific Ocean from USA, New Zealand, Australia, Korea between 2008 and 2012. Notably, one genome from Thau lagoon (MG-08-Me-Po-040) exhibited no admixture and represents a unique lineage associated with ancestral population 7 (Figure 2).

**Figure 1:**
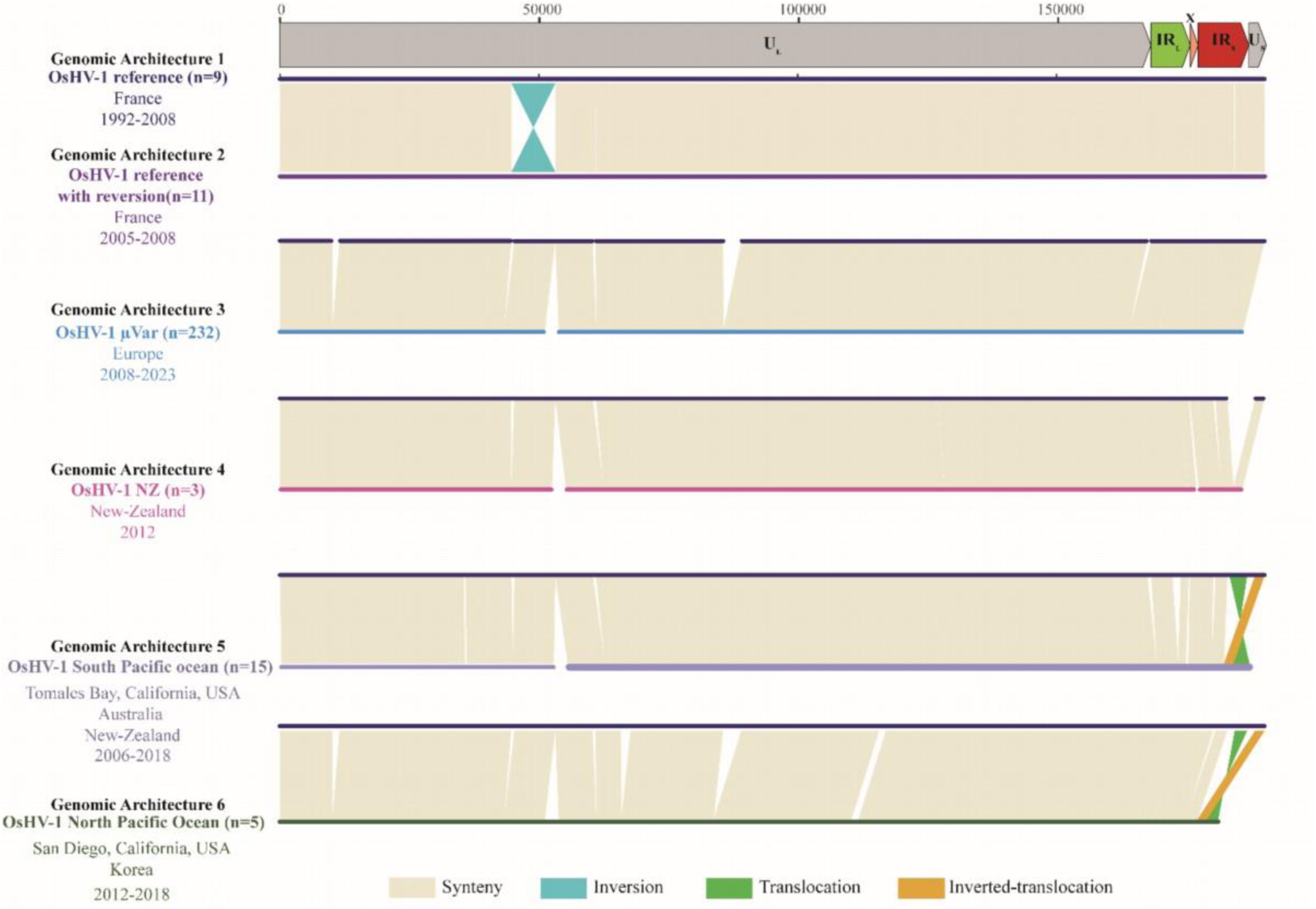
Comparative genomic showing the six OsHV-1 genomic structure type. The description of genomic structural variations was based on a pairwise comparison to the OsHV-1 reference genome (NC_005881.2, Davison et al. 2005) performed using Mummer. In each pairwise comparison, the reference genome is represented by a dark blue bar. Syntenic regions are represented in beige, inversions in light blue, translocations in green and inverted translocations in orange. Above the graph of structural variations is the architecture and position in the OsHV-1 reference genome.

**Figure 2:**
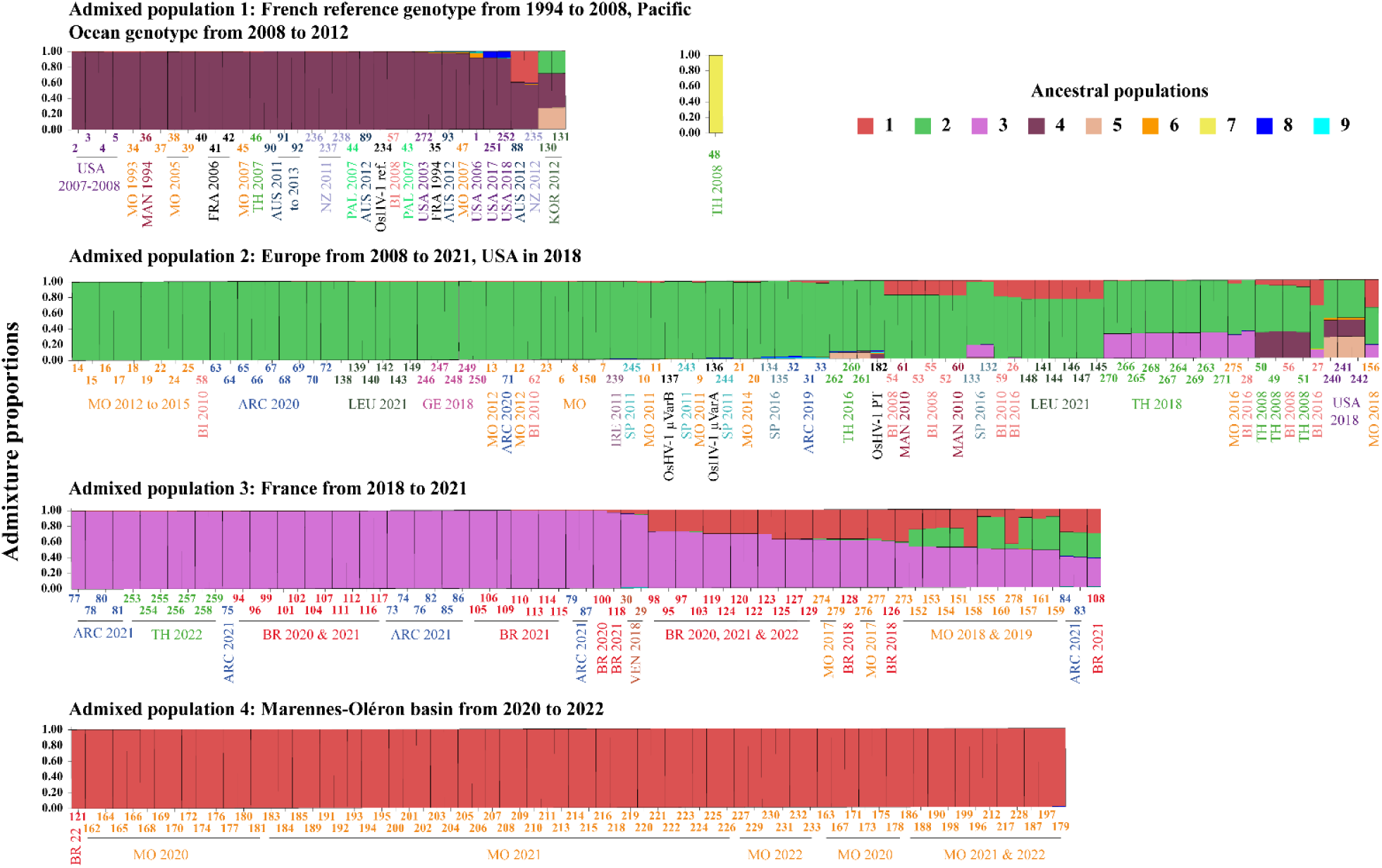
Population admixture and assignation of ancestral population to each of the 279 genomes. Bar plots representing the assignation of each individual OsHV-1 genome to an ancestral population based on admixture analysis. The analysis of population admixture has been performed using the STRUCTURE software, assuming nine ancestral populations (represented as red, green, pink, dark purple, salmon, orange, yellow, dark blue, light blue). The x axis represents OsHV-1 genomes from populations sorted according to heir reported ancestries (defined by Q-Matrix) and four groups of OsHV-1 genomes were defined arbitrary based on genome similarities regarding ancestral populations. Each individual OsHV-1 genome is epresented by a vertical stacked column of color-coded admixture proportions that reflects genetic contributions from putative ancestral populations.

The second admixed population includes genomes collected in Europe (Spain, Germany, Ireland, France) between 2008 and 2021, along with three genomes collected from San Diego in the USA in 2018. While predominantly associated with ancestral population 1, this group displays admixture from multiple populations, reflecting complex historical gene flow (Figure 2).

The third admixed population contains genomes collected exclusively in France between 2018 and 2021(Figure 2).

The last admixed population comprises genomes collected in the Marennes-Oléron basin from 2020 to 2022 and one genome from Brest harbor collected in 2022. These genomes are almost exclusively associated with a single ancestral population, indicating a recent, homogeneous lineage (Figure 2).

In France, five distinct patterns of population structure were observed. 169 genomes showed no admixture, 64 genomes displayed admixture between two or three ancestral populations (mainly populations 1, 2, and 3), and four lineages from Thau lagoon and Breton inlet collected in 2008 exhibited admixture involving populations 1, 2, and 4 (Figure 2).

In parallel, recombination events were explored to evaluate whether genetic exchanges between viral lineages could have contributed to the observed patterns of admixture. From the Gubbins recombination analysis (Croucher et al., 2015), 15 potential recombination events were identified along OsHV-1 genomes (Table S4), which might have impacted seven specific genomic regions (Figure S1). More specifically, these events were located around the stem loop (45,889 bp to 71,809 bp) and at the 3′ end of the U_L_ (152,920 bp to 167,829 bp), in a short fragment within the IR_L_ (172,129 bp to 172,233 bp) beginning of IR_L_, within the X region, at the end of IR_S_ (174,916 bp to 196,039 bp), and at the 3′ end of the U_S_ region (199,958 bp to 200,971 bp). Eleven potential recombination events occurred in the U_L_ region and five in the repeated regions (Figure S1). These regions were excluded from subsequent phylogenetic and population analyses to avoid bias.

### Network

A Median-Joining Network (MJN) was constructed from 279 NR-genome sequences to visualize the genetic relationships among OsHV-1 lineages worldwide (**Figure 3**). The resulting network revealed a clear structuring of viral diversity according to both geographic origin and sampling period. Three distinct clusters were observed: one comprising early European lineages sampled before 2008, a second encompassing strains from Oceania and the USA, and a third, more diverse cluster, containing numerous lineages sampled in Europe after 2008. Notably, sequences from Korea and the USA formed a small intermediate cluster linking early European and Pacific lineages. The largest and most reticulated part of the network corresponded to the post-2008 European lineages, characterized by a high number of mutational steps, frequent median vectors, and a dense network of derived lineages. This pattern suggests a recent and rapid diversification of the virus within Europe following its emergence in the late 2000s. In contrast, Pacific and early European lineages formed well-delineated and less reticulated clades.

**Figure 3:**
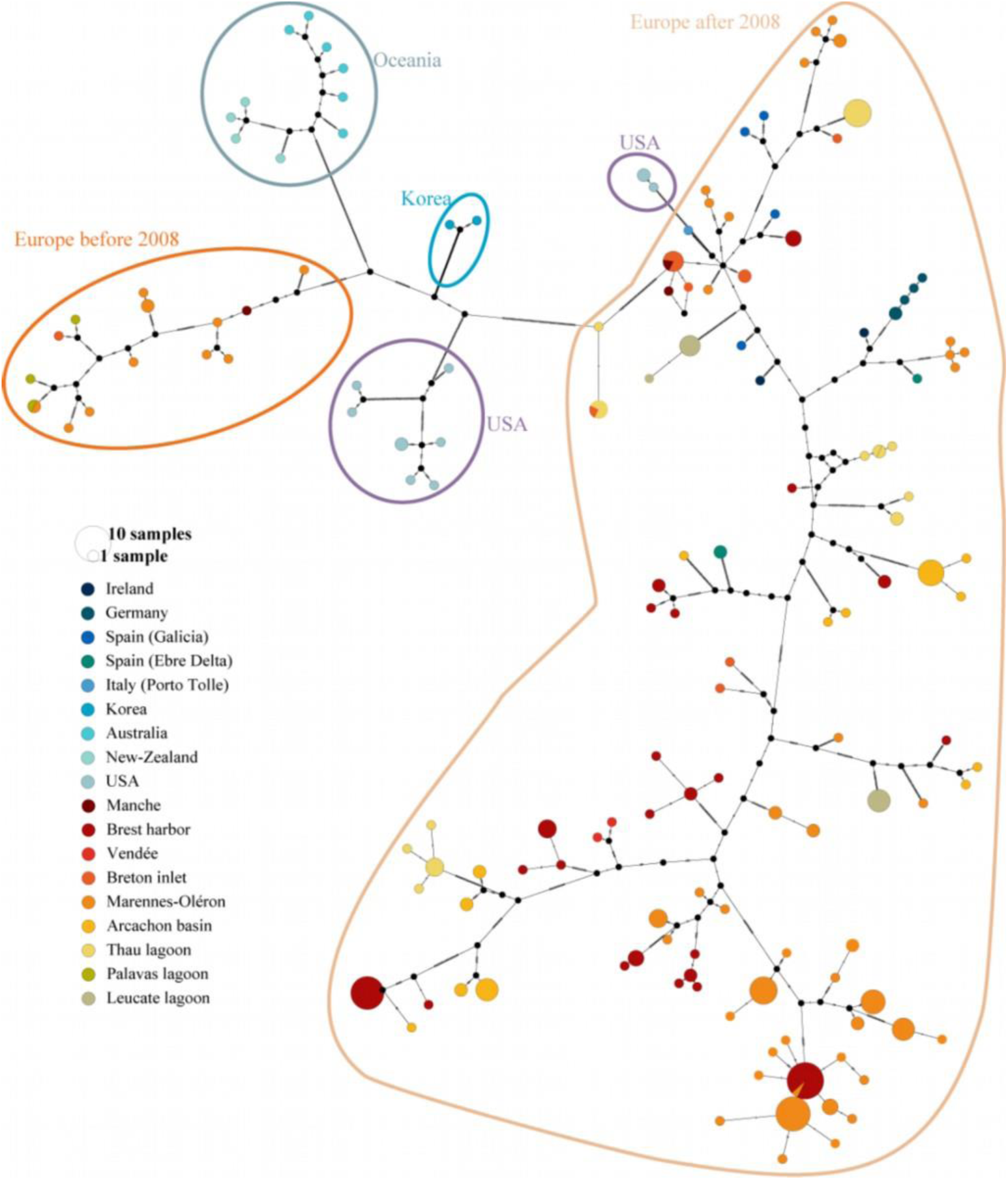
Median-Joining Network of 279 OsHV-1 genomes. Median-Joining Network (MJN) was generated from 279 OsHV-1 genomes (recombinant region excluded) to visualize global lineage relationships. Each node represents a unique lineage, with node size proportional to its frequency, and edges indicate mutational steps. Median vectors correspond to unsampled or ancestral lineages. The network reveals six major lineage groups: (i) two USA lineage groups in purple, (ii) Oceanian lineages in grey-blue, (iii) early European lineages sampled before 2008 in orange, (iv) Korean lineages in blue and (v) highly diverse European lineages sampled after 2008 in salmon.

### Maximum clade credibility tree and ancestral states inference

The regression of root-to-tip genetic distances against sampling dates (R² = 0.344) from the best-fitting root tree suggests OsHV-1 exhibits a clock-like evolutionary pattern. This indicates the presence of a sufficient temporal signal (*i.e.* meaningful correlation between genetic divergence and sampling time), allowing for the reliable application of molecular clock models to estimate evolutionary rates over time (**Figure 4**A).

**Figure 4:**
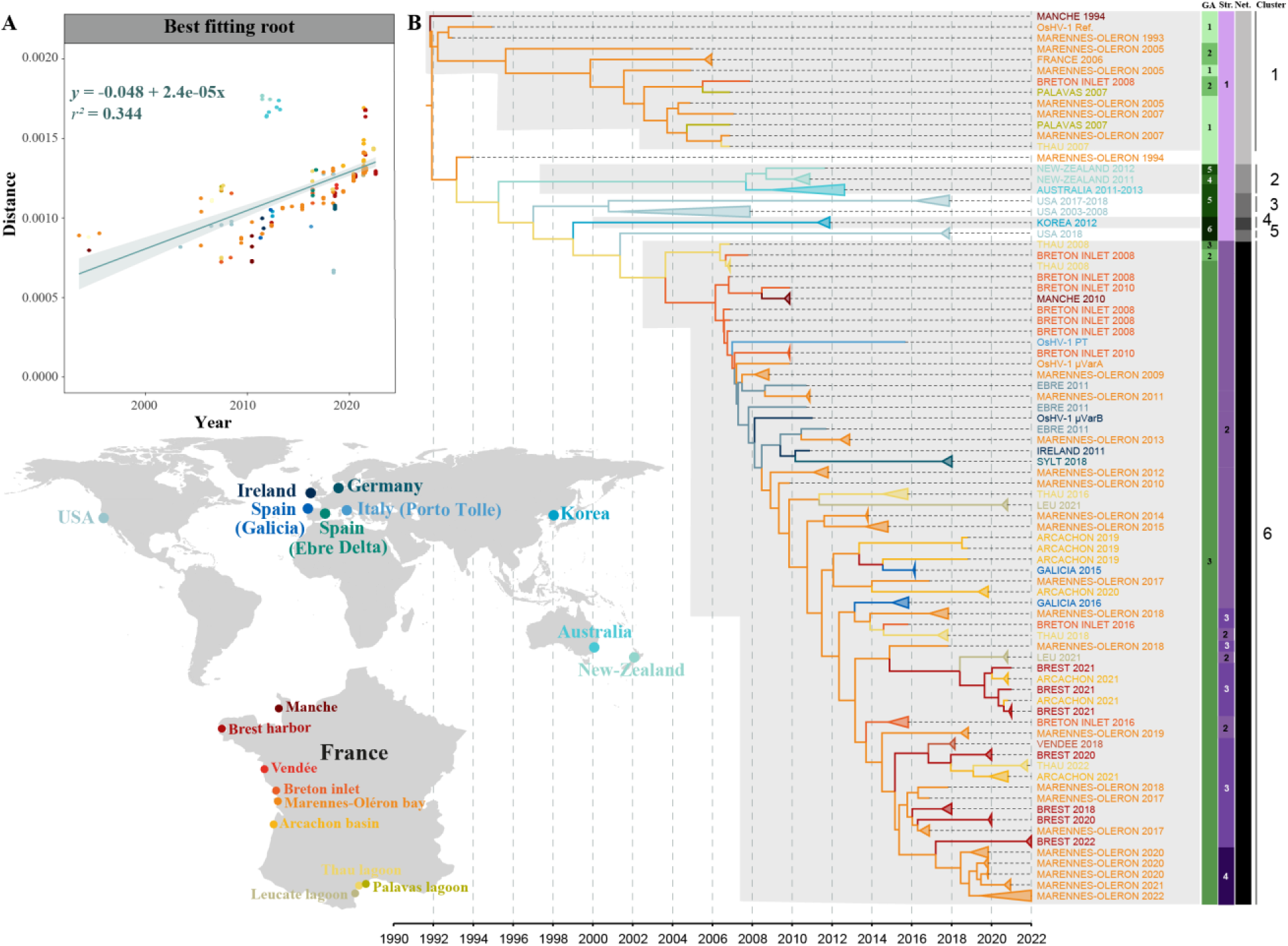
Ancestral state reconstruction of OsHV-1 specific origin between 1990 and 2022 worldwide. A) Root-to-tip distance in maximum likelihood tree versus sampling year. B) Time-scaled Maximum clade credibility of 279 OsHV-1 genomes which was obtained using Bayesian phylogenetic inference in program BEAST under a HKY+G+I model of nucleotide substitution, uncorrelated relaxed clock and GMRF demographical models. Tip colors correspond to origin of sampling and branch colors correspond to most probable origin of the common ancestor as specified on the map. This figure was produced in FigTree v1.4.4 (Rambaut, 2019). Six clusters were defined (numbered on the right).

The comparison of different discrete phylogeographic models of OsHV-1 dispersal showed statistical support for the combination of an uncorrelated relaxed molecular clock and a GMRF coalescent model with lower marginal likelihoods under path sampling (PS:-17810.75354) and stepping-stone sampling (SSS:-17820.08032) (Table S3).

The time-scaled phylogeny revealed six phylogenetic clusters (**Figure 4**B). The first phylogenetic cluster corresponds to lineages collected in France between 1993 and 2008, including the reference OsHV-1 lineage (Davison et al., 2005). The second cluster comprises OsHV-1 lineages collected on the South-West coasts of the Pacific Ocean (New-Zealand and Australia) between 2011 and 2013. Genomes collected from the North-East coasts of the Pacific Ocean (California, USA) between 2003 and 2018 formed a third phylogenetic cluster. Both OsHV-1 lineages collected from Korea in 2012 clustered in a fourth phylogenetic cluster. The fifth cluster grouped three OsHV-1 lineages collected from the USA in 2018. Finally, the last cluster encompassed lineages collected from European coasts (Germany, Italy, Spain, Ireland and France) between 2008 and 2022.

Furthermore, the mean substitution rate along the overall phylogeny was estimated to be 3.6093×10^-03^ nucleotide substitutions per site per year (ns.s^-1^.y^-1^) (95% HPD interval [3.2291×10^-03^ - 3.09444×10^-03^]). The most recent common ancestor (MRCA) of OsHV-1 sampled lineages was estimated to have existed in 1991 (95% HPD interval [1990-1992]) in the Marennes-Oléron basin (MRCA origin probability: 0.68). Subsequently, it can be inferred that this lineage might have evolved differently in Europe and the Pacific Ocean. In Europe, the first detected lineage was the OsHV-1 reference lineage sampled in 1992 (Davison et al., 2005). In parallel, from this reference lineage, another lineage might have emerged and could have been first detected in the Pacific Ocean between 1994 (95% HPD interval [1991-1998]) and 2018. Subsequently, in Europe, after the emergence of this lineage, it might have evolved into the OsHV-1 µVar lineage, which is currently the dominant lineage detected.

## Discussion

Understanding the dispersal and evolution of OsHV-1 is essential for managing oyster health and sustaining the global aquaculture industry, particularly given the virus’s severe impact on *M. gigas* populations. This study presents the first large-scale, spatio-temporal analysis of OsHV-1 genomic diversity restricted to isolates obtained from *M. gigas*, encompassing a wide geographic distribution and spanning three decades. By leveraging an unprecedented number of whole-genome sequences, this work integrates and builds upon previous research (Delmotte-Pelletier et al., 2022; Morga-Jacquot et al., 2021) and provides new insights into broad-scale patterns of viral diversification as well as fine-scale dispersal dynamics, with a particular focus on largest oyster-producing countries.

### From OsHV-1 to OsHV-1 µvar

The earliest OsHV-1 genomes analyzed in this study, spanning 1990–2008, correspond to the historical OsHV-1 reference lineage first described by Davison et al. (2005) and isolated in 1992. These genomes display the canonical 207 kb organization (TR_L_-U_L_-IR_L_-X-IR_S_-U_S_-TR_S_), occasionally accompanied by a ∼4 kb inversion within the U_L_ region. This inversion, found exclusively in a subset of French samples from 1993 to 2008, was previously documented in a minority (25 %) of the population, affects ORFs 36-43 and disrupts ORF 36, likely rendering the encoded membrane protein non-functional (Davison et al., 2005). The remaining ORFs, despite being inverted, remain intact, and any functional impact on them may be reversible. Such structural rearrangements are common in herpesviruses and may generate defective viral genomes requiring complementation by intact viral particles to complete the replication cycle (Vignuzzi & López, 2019).

Around 2008, the genomic architecture of OsHV-1 circulating in France and Europe shifted markedly with the emergence of the OsHV-1 µVar lineage, signifying “microsatellite variation” (Segarra et al., 2010). Extensive retrospective analyses of archived samples collected before 2008 did not detect the µVar lineage, supporting its absence from European oyster populations prior to its first description. This lineage is characterized by a genome of 204 kb and defined by a constellation of structural variations totaling nearly 10 kb, including: (*i*) a characteristic 12-bp deletion in the microsatellite upstream of ORF 4, encoding a protein of unknown function, (*ii*) an insertion of ∼2.7 kbp between ORF 43 (encoding a protein of unknown function) and ORF 44 (encodes a Deoxynucleoside kinase domain), *(iii)* five large deletions spanning the U_L_ and IR_L_ regions for a total of ∼7 kbp. OsHV-1 µVar rapidly replaced the reference lineage in Europe, after a first detection in France and became predominant after 2008, a transition associated with increased oyster mortalities (Burioli et al., 2017; Segarra et al., 2010).

Population genetic and recombination analyses support a more complex evolutionary origin of OsHV-1 µVar than previously appreciated. Four genomes collected in 2008 from Thau Lagoon and the Breton Inlet exhibit genetic admixture between µVar and the ancestral reference lineage. Recombination signals were detected in up to eight genomes from that year, involving regions surrounding the stem-loop structure, the ORF 4 microsatellite, and multiple ORFs encoding transmembrane or zinc-finger proteins. The simultaneous detection of admixed genomes in two geographically distant French production areas - despite likely no natural hydrodynamic connectivity - suggests a scenario in which µVar emerged in one basin and was subsequently disseminated through oyster transfers (Keeling et al., 2001; Lupo et al., 2016; Morse, 2004). However, limited sampling during 2008 and the absence of precise metadata for several genomes preclude definitive conclusions. Finally, it’s worth noting that the sampling was exclusively conducted within these two basins, which might not provide a comprehensive representation of the broader context.

These results support a scenario in which the µVar lineage did not arise from a gradual accumulation of point mutations but from a recombination-driven restructuring of the reference genome, likely facilitated by intense oyster movements, high farming densities, and the presence of mixed viral populations within hosts. This evolutionary shift may also have been reinforced by periods of strong natural recruitment and the use of hatchery-produced spat, which together contributed to very high densities of juvenile oysters. Such high densities of susceptible hosts may have favored intense viral circulation and increased viral pressure, potentially facilitating the emergence of new variants (Geay & Mille, 2008). Such demographic amplification would have enhanced viral transmission opportunities and co-infection events, thereby promoting recombination and accelerating the emergence of OsHV-1 µVar

### The emergence and evolution of Pacific OsHV-1 lineages

Between 2006 and 2018, OsHV-1 lineages sampled in Pacific regions, including the USA (Tomales Bay and San Diego), Australia, New Zealand, and Korea, revealed distinct genomic architectures that differ from both the OsHV-1 reference and the European OsHV-1 µVar lineages. These Pacific lineages exhibit extensive rearrangements and deletions in the U_L_ region, notably affecting ORF 4 and ORFs 42/43, while retaining elements absent from European µVar, such as intact ORFs 62/63 and partial microsatellite sequences upstream of ORF 4. Short tandem repeats are common and highly polymorphic in non-coding regions of herpesvirus genomes, including HSV-1, where untranslated regions can regulate viral gene expression (Bagshaw, 2017; Deback et al., 2009). Although their function remains unclear, microsatellite instability may generate frameshift mutations and phenotypic variation that could influence virulence or host immune gene escape (Bagshaw, 2017; Deback et al., 2009; Ramchandani et al., 2019; Srivastava et al., 2019; Wu et al., 2014).

Newly identified large-scale rearrangements, including translocation of the U_S_ region and reverse translocation of the IR_L_ region, further distinguish these Pacific lineages from the others sampled. These structural configurations are consistent with an evolutionary trajectory that unfolded outside Europe and likely reflects long-term diversification within the broader Indo-Pacific region. Previous molecular detection first confirmed the presence of OsHV-1 outside Europe on the west coast of North America in 2002 (Friedman et al., 2005), where lineages related to the OsHV-1 reference were identified (Grijalva-Chon et al., 2013; Renault et al., 2012). By 2010, OsHV-1 µVar was detected in Australia and New Zealand (Jenkins et al., 2013), revealing novel lineages that, while related to previously described lineages, displayed distinct molecular signatures. More recently, full-genome characterizations of Australian lineages (Trancart et al., 2022) confirmed the presence of substantial structural rearrangements, supporting the idea that Pacific viral populations have accumulated region-specific genomic architectures.

Population admixture, network and phylogeographic analyses position these Pacific lineages intermediate between the OsHV-1 reference lineage, and the European µVar lineage, suggesting that recombination among these ancestral populations contributed to the emergence of the Pacific lineages. This pattern is coherent with the extensive viral diversity documented in East Asia. A first large-scale molecular survey based on marker ‘A’ within U_L_ region, upstream of ORF 83 (Renault et al., 2000) revealed substantial variation across multiple *Crassostrea* species distributed from subtropical to temperate regions of China, Korea, and Japan (Moss et al., 2007). Subsequent analyses using the more informative marker ‘C’ (located within the TR_L_/IR_L_ regions, in a non-coding region, and a fragment of ORF4, Batista et al., 2015), uncovered an even broader genotypic spectrum in the Japanese archipelago, including numerous unique lineages detected in asymptomatic *M. gigas* and related species (Shimahara et al., 2012). Additional sequences from China and Korea further highlighted the presence of distinctive and sometimes closely related lineages within the Northwest Pacific (Hwang et al., 2013; Renault et al., 2012). Together, these data indicate that East Asia constitutes a major reservoir of OsHV-1 diversity, within which recombination and structural reshuffling likely occurred over extended evolutionary timescales.

Recurrent deletions represent a hallmark of Pacific OsHV-1 evolution, affecting ORFs 35–37, ORF 48, and ORFs 62–63, with ORF 36 frequently disrupted or absent in several lineages. The recurrence of deletions and inversions of ORF 36 across geographically distant Pacific regions suggests that this locus may represent a conserved hotspot for structural evolution. Such patterns, observed in North American, New-Zealand and Australian genomes as well as in East Asian lineages, indicate that genomic plasticity is a pervasive and intrinsic feature of OsHV-1 evolution outside Europe. These deletions, together with large-scale rearrangements, likely enhance viral adaptability by increasing genomic variability, thereby facilitating persistence across diverse climatic regimes and host assemblages. The generation of these structural variations is plausibly driven by herpesvirus-specific mechanisms, including homologous recombination, genome rearrangements mediated by repeated regions, and the formation of alternative genomic isomers during replication. The high nucleotide diversity measured across the Northwest Pacific, combined with the broad host range encompassing *M. gigas*, *C. ariakensis*, *C. sikamea*, and other sympatric species (Moss et al., 2007; Shimahara et al., 2012), suggests that long-standing host–virus coevolutionary processes have shaped viral genomic architecture in this region. Rather than reducing viral fitness, these structural dynamics may confer selective advantages by promoting lineage diversification and ecological persistence.

### Evolutionary forces shaping the global diversity of OsHV-1

The estimated mean evolutionary rate of OsHV-1 (3.6 × 10^⁻03^ nucleotide substitutions per site per year) falls between previous estimates derived from smaller datasets and different estimation methods (Morga-Jacquot et al., 2021; Pelletier et al., 2025). Although methodological differences partly explain discrepancies among studies, this rate remains higher than those reported for several other herpesviruses, including HSV-1 (8.21×10^⁻05^ ns.s^-1^.y^-1^), VZV (6.26×10^-06^ ns.s^-1^.y^-1^) and Marek’s disease virus (∼1.6×10^⁻05^ ns.s^-1^.y^-1^) (Sanjuán, 2012; Sanjuán et al., 2010; Trimpert et al., 2017). Together with the numerous insertions, deletions, and recombination signals detected across genomes in this study, these results indicate that multiple evolutionary processes contribute to OsHV-1 diversification. Such comparatively elevated rates suggest that OsHV-1 may experience strong or frequent evolutionary pressures. While, uneven sampling across countries - with disproportionate genome availability from specific areas such as Marennes-Oléron - can influence molecular clock estimates and inflate apparent rates (Gámbaro et al., 2025); several other non-exclusive factors may contribute to this pattern.

First, high host densities in intensive aquaculture systems enhance viral transmission and replication opportunities, thereby increasing the probability of mutation accumulation (Kennedy et al., 2016; Pernet et al., 2016). Second, ecological overlap between oysters and other mollusks and occasional multi-species farming systems may facilitate interspecific transmission and recombination events, processes already documented in OsHV-1 genomes and widely recognized in herpesvirus evolution (Casto et al., 2020; Crespo-Bellido et al., 2021). In addition to nucleotide substitutions, recurrent large insertions and deletions affecting several ORFs and regulatory regions highlight the importance of structural variation as a driver of OsHV-1 genomic plasticity. Recombination signals detected among co-circulating lineages further suggest that coinfections within hosts can generate mosaic genomes, accelerating diversification beyond what would be expected from point mutations alone.

Ecological and aquaculture dynamics likely played a central role in shaping Pacific OsHV-1 diversity. The global expansion of *M. gigas* from its native Northwest Pacific range, facilitated by its adaptability and aquaculture value (Andrews, 1980), created repeated opportunities for viral dissemination across ocean basins. Intensification of farming practices, large-scale spat exchanges between hatcheries and production areas, and the maintenance of high oyster densities have likely amplified viral circulation. Importantly, oysters may acquire infection during early developmental stages without showing clinical signs (Dégremont, 2013; Peeler et al., 2012), enabling asymptomatic carriage, as observed in both wild and farmed oysters across East Asia, North America, Australia, and New Zealand, which can facilitate inadvertent viral spread during stock movements. These processes provide a plausible ecological framework for the persistence, diversification, and regional structuring of OsHV-1 lineages.

### Conclusions and perspectives

Altogether, the integration of genomic architecture, time-scaled phylogenies and population structure analyses reveals a complex and dynamic evolutionary landscape of OsHV-1 at both regional and global scales. Our results highlight long-standing viral diversity in East Asia, the emergence of structurally distinct Pacific and microvariants lineages, and ongoing diversification driven by the combined effects of nucleotide substitutions, recurrent insertions and deletions, recombination, and large-scale genomic rearrangements, probably reinforced by anthropogenic oyster movements. By combining three decades of whole-genome data from major oyster-producing regions worldwide, this study provides a robust baseline for understanding OsHV-1 diversity, evolution, and transmission dynamics. Our findings further support a scenario in which the emergence of the µVar lineage was driven primarily by genomic restructuring rather than by the gradual accumulation of point mutations, a process likely facilitated by intensive oyster transfers, high farming densities, and the expansion of hatchery-produced spat that increased host connectivity among farming areas. However, the present data do not allow to definitively determine whether the µVar lineage emerged locally in France or was introduced from another country through oyster farming practices and associated transfers. These insights are critical for improving molecular surveillance, anticipating viral spread, and developing targeted, regionally adapted management strategies aimed at mitigating the impact of OsHV-1 on oyster populations and ensuring the long-term sustainability of global aquaculture systems. Future work should aim to identify the ecological, environmental, and husbandry-related factors shaping viral diversity, including temperature regimes, host density, and farming practices, in order to better disentangle their respective contributions to viral evolution. Expanding genomic sampling to underrepresented regions, particularly in Asia, Oceania, and emerging aquaculture areas, will be essential to capture the full extent of global diversity and refine phylogeographic inferences. Integrating these approaches with longitudinal sampling and epidemiological data will further improve our ability to track transmission pathways and predict the emergence of novel variants.

## Material and methods

### Samples collection

A total of 275 naturally infected *M. gigas* were collected between 1993 and 2022 (Table S1, **Figure 5 Erreur ! Source du renvoi introuvable.**). These samples originated from different country around the world (Figure 1A): Ireland (n=1), Korea (n=2), New-Zealand (n=4), Germany (n=5), Australia (n=6), Spain (n=7), USA (n=11) and France (n=238). In France, samples were collected at nine different locations (Figure 1B): Palavas lagoon (Hérault, n=2), in Bouin (Vendée, n=2), in the Manche channel (Normandie, n=3), in the Leucate lagoon (Aude, n=12), in the Breton inlet (Charente-Maritime and Vendée, n=12), in the Thau lagoon (n=24), in Arcachon basin (Gironde, n=28), in the Brest harbor (Finistère, n=38), and in the Marennes-Oléron bay (Charente-Maritime, n=113). Additionally, four samples were collected in France for which the exact location is unknown.

**Figure 5:**
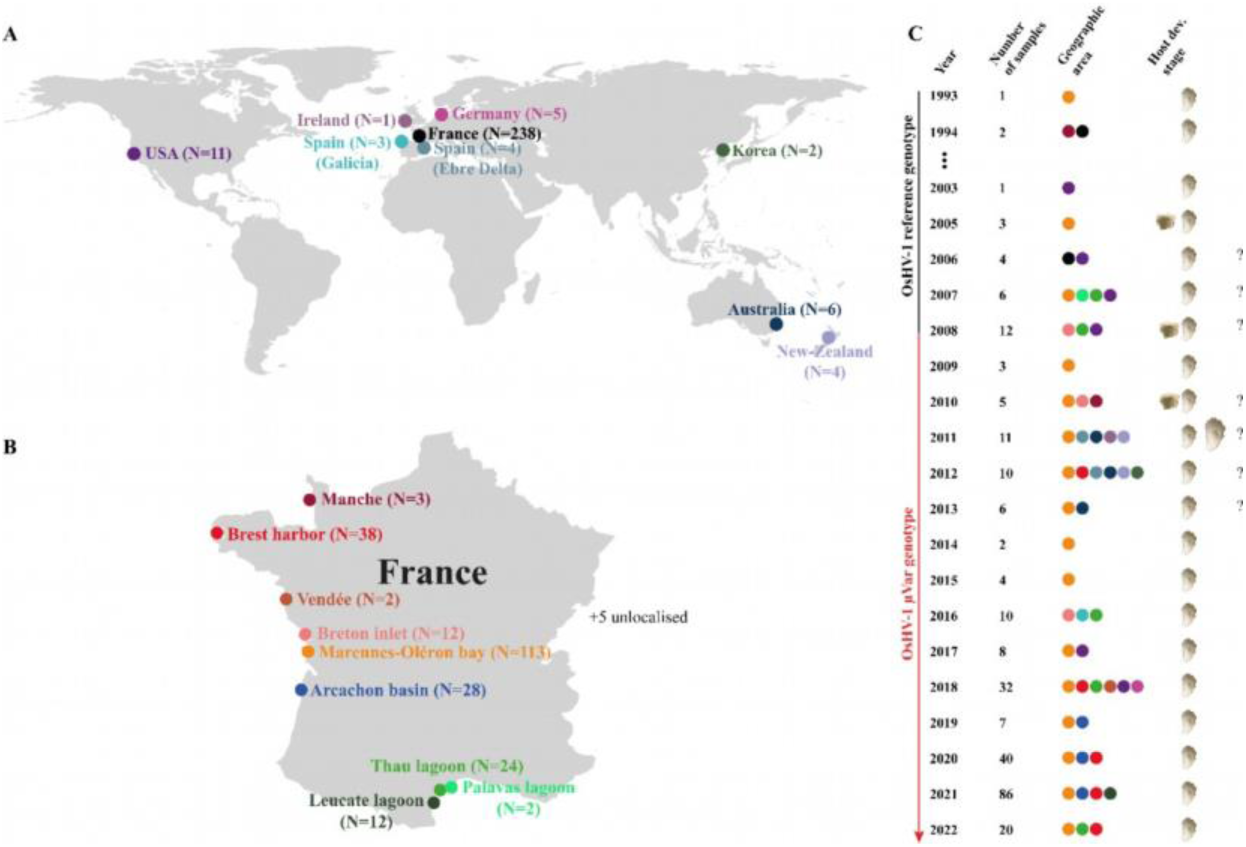
Samples origin and sampling metadata. A) Overview of locations and number of samples collected around the world. B) Overview of locations and number of samples collected in France. C) Year of sampling and number of samples, geographic areas and host developmental stages associated.

### DNA extraction, viral copy numbers quantification and sequencing

DNA was extracted from the mantle tissue of individual moribund oysters using either the MagAttract® HMW DNA kit or QIAamp DNA mini kit (Qiagen) according to manufacturer’s protocols (Table S1). DNA purity and concentration were measured with a Nano-Drop ND-1000 spectrophotometer (Thermo Scientific) and quantified using Qubit^®^ dsDNA BR assay kits (Molecular Probes Life Technologies). Viral load was determined by quantitative PCR using a Mx3005 P thermocycler (Agilent), following the method described in Pepin et al. (2008).

Illumina DNA-seq library were prepared and sequenced across six different sequencing platforms (Table S1). One sample was sequenced at Genotoul (INRAE, Castanet-Tolosan, France), three samples were sequenced at GATC biotech (Eurofins Scientific, Eberberg, Germany), 26 samples were sequenced at LIGAN-PM Genomics (CNRS UMR 8199, Lille, France), 161 samples were sequenced at the Genome Quebec Company (Genome Quebec Innovation Center, McGill University, Montreal, Canada), 24 samples were sequenced at Fasteris Life Science Genesupport SA (Plan-les-Ouates, Switzerland) and 59 samples were sequenced at IntegraGen (Evry, France).

### *De novo* assembly of OsHV-1 genomes

The assembly of viral genomes is essential for investigating viral diversity, as it enables the reconstruction of complete genomes from infected samples. The *de novo* assembly of OsHV-1 genomes were reconstructed using the analytical framework previously developed (Delmotte-Pelletier et al. 2022 and Dotto-Maurel et al. 2022). Briefly, Illumina reads were quality-filtered and trimmed with Fastp (version 0.20.1, Chen et al., 2018) before removing host-derived sequences by mapping against the *M. gigas* genome (accession: GCA_902806645.1) using Bowtie2 (version 2.4.1, Langmead & Salzberg, 2012). Non-oyster reads were assembled *de novo* with SPAdes (version 3.14.0, with parameter –meta, Bankevich et al., 2012), and OsHV-1 contigs were identified by BLASTn (version 2.6.0, Altschul et al., 1990) and manually scaffolded in Geneious Prime (version 2020.2.3) to reconstruct the canonical U_L_–IR_L_–X–IR_S_–U_S_ architecture. Because repeated regions cannot be reliably resolved with short reads, we generated non-redundant genomes (NR-genomes) containing a single copy of each repeat.

To ensure the accuracy and consistency of the final assemblies, quality-filtered reads were realigned to the newly reconstructed OsHV-1 NR-genomes using Bowtie 2 v2.4.1 (parameter --end-to-end). Raw data and reconstructed genomes have been deposited in the ENA database for future reference and accessibility (Table S1).

### Whole genome alignment and recombination detection

#### Identification and description of OsHV-1 structural variations

At the genomic scale, variations between different lineages of OsHV-1 can be attributed to structural variations (SV) such as insertions, deletions, inversions, translocations, and duplications. To analyze and compare the genomic architecture among viral lineages and discern large-scale evolutionary events, we aligned the 274 reconstructed NR-genomes with four published OsHV-1 genomes using the progressiveMauve algorithm implemented in Mauve v2.4.0 with default parameters (Darling et al., 2004). These four genomes were selected because they were also isolated from the *M. gigas* and include OsHV-1 PT (2016, Italy: MG561751; Abbadi et al., 2018), OsHV-1 µVarA (2010, France: KY242785; Burioli et al., 2017), OsHV-1 µVarB (2011, Ireland: KY271630; Burioli et al., 2017), and OsHV-1 Ref. (1992-1993, France: NC_005881.2; Davison et al., 2005).

Pairwise and multiple whole-genome alignments of these representative lineages against the OsHV-1 reference genome (NC_005881.2) were performed using the progressiveMauve algorithm implemented in Mauve v2.4.0 with default parameters (Darling et al., 2004) to visualize structural variations.

#### OsHV-1 genomes in silico final polishing and multiple alignment

To conduct the phylogenetic analysis, it was essential to standardize genomic structures across all genomes. Because 34 genomes displayed substantial rearrangements relative to OsHV-1 reference genome (Davison et al., 2005), manual adjustments were made to these assemblies. Using Mauve multiple alignment in Geneious Prime, the genomic architecture of each genome was carefully standardized, ensuring consistency and enabling meaningful comparisons. The genomes were then aligned using MAFFT v1.4.0 (Katoh et al., 2002).

#### OsHV-1 populations admixture

In order to assess the potential influence of geographic origin on the structure of the OsHV-1 population, a population structure analysis was conducted. First, Single Nucleotide Polymorphisms (SNP) were obtained from the multiple alignment of all 278 NR-genomes previously generated and extracted using SNP-sites v2.5.1 (Page et al., 2016). Next, we used the software STRUCTURE v 2.3.4 to estimate population admixture and to assign each individual to ancestral populations (Pritchard et al., 2000). To determine the optimal number of populations (K), independent runs of the Gibbs sampler (Robert & Casella, 1999) were executed with K values ranging from 1 to 20. The Gibbs sampler applied a Bayesian approach using Markov Chain Monte Carlo inference. Each run consisted of 50,000 iterations, with a burn-in period of 5,000 iterations. The STRUCTURE software assigned a posterior probability to each tested population structure, which was used to select the appropriate number of populations as described by Porras-Hurtado et al. (2013)(Table S3). Based on this analysis, K=9 was selected to infer population structure. To refine the population structure, a final run with 100,000 iterations, and a burn-in period of 10,000 iterations was performed for K=9. Finally, bar plots representing the estimated membership coefficients were generated with STRUCTURE using the output Q-matrix and groups (*i.e.* assignment to pre-defined populations) of individuals were defined arbitrarily based on ancestral similarities.

#### Recombination events detection

The presence of recombinant regions along alignments can introduce biases in tree reconstruction due to the potentially distinct evolutionary histories of those regions compared to the rest of the genome (Lanier & Knowles, 2012). Beyond their methodological impact, however, identifying such regions is essential for understanding the evolutionary mechanisms shaping pathogen diversification, as recombination constitutes a major driver of genetic innovation and adaptation (Pérez-Losada et al., 2015; Simon-Loriere & Holmes, 2011). To address this issue, we employed Gubbins v3.3 (Croucher et al., 2015) with default settings to detect regions in the assembled OsHV-1 NR-genomes that may have been affected by recombination events. Any genomic regions identified as potentially recombinant were subsequently excluded from the downstream analyses (Table S2).

#### Genetic network construction

A Median-Joining Network (MJN) was constructed to visualize genetic relationships among sequenced genomes. A multiple alignment of all 279 NR-genomes without evidence of recombination was used as input in PopART v1.7 (Leigh & Bryant, 2015). Prior to network construction, all ambiguous sites and missing data were removed to minimize reconstruction artifacts. The MJN algorithm was applied with default parameters, and epsilon was set to zero to reduce the complexity of the resulting network. Each unique genome was represented as a node, with node size proportional to its frequency in the dataset. Mutational steps between haplotypes were represented by connecting lines, and median vectors were automatically inferred to account for unsampled or ancestral sequences. The resulting network was used to identify patterns of genetic structure and potential relationships among sampled individuals.

#### Phylogeographic inferences

To reconstruct the worldwide ancestral origin of OsHV-1 a discrete asymmetric phylogenetic diffusion model was used in BEAST v1.10.4 (Drummond & Rambaut, 2007). This involved modeling the host origin country as a discrete trait for each NR-genome sequence over the phylogenetic tree and inferring ancestral states. To estimate the likelihood of internal nodes and branches being associated with a specific origin, ancestral state inference based on host information from the branch tips was employed. The best fitting model of evolution for the samples was found using MEGA v11.0.11 (Tamura et al., 2021) based on Akaike Information Criterion (AIC). Our analysis used a HKY+G+I model of nucleotide substitution.

To identify the most suitable combination of coalescent (*i.e.* constant size, Gaussian Markov Random Field or GMRF) and molecular clock models (*i.e.* strict clock and uncorrelated relaxed clock) for our dataset, each possible combination has been tested. The maximum likelihood estimator (MLE) was calculated using path sampling (PS) and stepping-stone sampling (SSS) for each model. Based on these estimators, an uncorrelated relaxed molecular clock and a GMRF coalescent model was defined as most appropriate. A Markov Chain Monte Carlo (MCMC) with 10,000,000 steps, sampling every 1,000,000 generations using Beast v1.10.4 was ran. Additionally, a Markov jumps approach (Minin & Suchard, 2008a, 2008b) was used to determine the number of transitions between spatial locations. The BEAGLE library was used to increase computational speed (Ayres et al., 2012).

To assess the mixing of the population (Ne > 200) and the stationary distribution of the trace, we examined the effective sample size in Tracer v1.7.2 (Rambaut et al., 2018). This control was performed for all parameters, with an initial burn-in of 10 %. Finally, we generated maximum clade credibility (MCC) trees using the tree output files from BEAST and annotated them in FigTree v1.4.4 (http://tree.bio.ed.ac.uk/software/figtree/).

## Data availability

The datasets generated from this study can be found in the SRA database BioProject (Table S1). All code is available on GitHub at the following URL https://gitlab.ifremer.fr/asim/oshv-1-phylogeography/-/tree/c08d8196deba9a6a1d3d3747d13b378eb23f2ad0/.

Complementary information is available from the corresponding authors upon reasonable request.

## Acknowledgement

This work was supported by the French Institute for Exploitation of the Sea (Ifremer), by DGAL (French General Directorate for Food) through the National Reference Laboratory for Mollusc Diseases and by the European-Union Reference Laboratory for Mollusc Diseases, Ifremer, La Tremblade. Additionally, it was supported by the EU project VIVALDI (H2020 program, no. 678589) led by Ifremer. We sincerely thank the French National Research Agency (ANR) for the time and support dedicated to the development of this work (IDEAL project: ANR-23-CE35-0009). CP was supported by grant from the region Nouvelle-Aquitaine, from the scientific direction of Ifremer and the ANR-23-CE35-0009 project.

The study was conceptualized and designed by BM and CP. BM and CP were responsible for sample collection and conducting experimental procedures. CP, GC and MJ performed bioinformatics and statistical analyses. CP drafted the manuscript, with all authors contributing to its revision and approval.

The authors would like to express their gratitude to the staff at the Ifremer stations in Argenton (LPI, PFOM) and La Tremblade (PMMLT) for their technical support in producing standard (NSI) oysters and conducting transplantation experiments. Special thanks go to Nicole Faury (Ifremer, ASIM) for their technical assistance. Thank you to Jean-Michel Escoubas (IHPE, Ifremer, Montpellier), Bruno Petton (LEMAR, Ifremer, Argenton), and Johan Vieira (Capena, Arcachon) for their assistance during field monitoring activities. Thanks to all the worldwide collaborators: Colleen Burge (Coastal and Marine Sciences Institute, California), Mathias Wegner (AWI, Germany), Stein Mortensen (Institute of Marine Research, Norway), Yoshiko Shimahara (National Reasearch Institute of Aquaculture, Fisheries Research Agency, Japan).

## References

Abbadi, M., Zamperin, G., Gastaldelli, M., Pascoli, F., Rosani, U., Milani, A., Schivo, A., Rossetti, E., Turolla, E., Gennari, L., Toffan, A., Arcangeli, G., & Venier, P. (2018). Identification of a newly described OsHV-1 µvar from the North Adriatic Sea (Italy). Journal of General Virology, 99(5), Article 5. 10.1099/jgv.0.001042

Altschul, S. F., Gish, W., Miller, W., Myers, E. W., & Lipman, D. J. (1990). Basic local alignment search tool. Journal of Molecular Biology, 215(3), Article 3. 10.1016/S0022-2836(05)80360-2

Andrews, J. D. (1980). A Review Of Introductions Of Exotic Oysters And Biological Planning For New Importations.

Arzul, I., Corbeil, S., Morga, B., & Renault, T. (2017). Viruses infecting marine molluscs. Journal of Invertebrate Pathology, 147, 118–135. 10.1016/j.jip.2017.01.009

Arzul, I., Nicolas, J.-L., Davison, A. J., & Renault, T. (2001). French Scallops: A New Host for Ostreid Herpesvirus-1. Virology, 290(2), Article 2. 10.1006/viro.2001.1186

Arzul, I., Renault, T., Lipart, C., & Davison, A. J. (2001). Evidence for interspecies transmission of oyster herpesvirus in marine bivalves. Journal of General Virology, 82(4), Article 4. 10.1099/0022-1317-82-4-865

Bagshaw, A. T. M. (2017). Functional Mechanisms of Microsatellite DNA in Eukaryotic Genomes. Genome Biology and Evolution, 9(9), 2428–2443. 10.1093/gbe/evx164

Bai, C.-M., Morga, B., Rosani, U., Shi, J., Li, C., Xin, L.-S., & Wang, C.-M. (2019). Long-range PCR and high-throughput sequencing of Ostreid herpesvirus 1 indicate high genetic diversity and complex evolution process. Virology, 526, 81–90. 10.1016/j.virol.2018.09.026

Bankevich, A., Nurk, S., Antipov, D., Gurevich, A. A., Dvorkin, M., Kulikov, A. S., Lesin, V. M., Nikolenko, S. I., Pham, S., Prjibelski, A. D., Pyshkin, A. V., Sirotkin, A. V., Vyahhi, N., Tesler, G., Alekseyev, M. A., & Pevzner, P. A. (2012). SPAdes: A New Genome Assembly Algorithm and Its Applications to Single-Cell Sequencing. Journal of Computational Biology, 19(5), Article 5. 10.1089/cmb.2012.0021

Batista, F. M., López-Sanmartín, M., Grade, A., Morgado, I., Valente, M., Navas, J. I., Power, D. M., & Ruano, F. (2015). Sequence variation in ostreid herpesvirus 1 microvar isolates detected in dying and asymptomatic Crassostrea angulata adults in the Iberian Peninsula: Insights into viral origin and spread. Aquaculture, 435, 43–51. 10.1016/j.aquaculture.2014.09.016

Battistel, C., Mouren, J.-C., Morga, B., Pelletier, C., Canier, L., Garcia, C., Arzul, I., Pihan, Y., Leroi, L., Chevignon, G., Durand, P. G., & Jacquot, M. (2025). MoPSeq-DB: A user-friendly web application for genomic data management and analysis of marine mollusc pathogens. Database, 2025, baaf080. 10.1093/database/baaf080

Brown, J. W., & Smith, S. A. (2018). The Past Sure is Tense: On Interpreting Phylogenetic Divergence Time Estimates. Systematic Biology, 67(2), 340–353. 10.1093/sysbio/syx074

Burioli, E. A. V., Prearo, M., & Houssin, M. (2017). Complete genome sequence of Ostreid herpesvirus type 1 µVar isolated during mortality events in the Pacific oyster Crassostrea gigas in France and Ireland. Virology, 509, 239–251. 10.1016/j.virol.2017.06.027

Casto, A. M., Roychoudhury, P., Xie, H., Selke, S., Perchetti, G. A., Wofford, H., Huang, M.-L., Verjans, G. M. G. M., Gottlieb, G. S., Wald, A., Jerome, K. R., Koelle, D. M., Johnston, C., & Greninger, A. L. (2020). Large, Stable, Contemporary Interspecies Recombination Events in Circulating Human Herpes Simplex Viruses. The Journal of Infectious Diseases, 221(8), 1271–1279. 10.1093/infdis/jiz199

Chen, S., Zhou, Y., Chen, Y., & Gu, J. (2018). fastp: An ultra-fast all-in-one FASTQ preprocessor. Bioinformatics, 34(17), Article 17. 10.1093/bioinformatics/bty560

Crespo-Bellido, A., Hoyer, J. S., Dubey, D., Jeannot, R. B., & Duffy, S. (2021). Interspecies Recombination Has Driven the Macroevolution of Cassava Mosaic Begomoviruses. Journal of Virology, 95(17), e00541–21. 10.1128/JVI.00541-21

Croucher, N. J., Page, A. J., Connor, T. R., Delaney, A. J., Keane, J. A., Bentley, S. D., Parkhill, J., & Harris, S. R. (2015). Rapid phylogenetic analysis of large samples of recombinant bacterial whole genome sequences using Gubbins. Nucleic Acids Research, 43(3), e15. 10.1093/nar/gku1196

Darling, A. C. E., Mau, B., Blattner, F. R., & Perna, N. T. (2004). Mauve: Multiple Alignment of Conserved Genomic Sequence With Rearrangements. Genome Research, 14(7), 1394–1403. 10.1101/gr.2289704

Davison, A. J., Trus, B. L., Cheng, N., Steven, A. C., Watson, M. S., Cunningham, C., Deuff, R.-M. L., & Renault, T. (2005). A novel class of herpesvirus with bivalve hosts. Journal of General Virology, 86(1), Article 1. 10.1099/vir.0.80382-0

Deback, C., Boutolleau, D., Depienne, C., Luyt, C. E., Bonnafous, P., Gautheret-Dejean, A., Garrigue, I., & Agut, H. (2009). Utilization of Microsatellite Polymorphism for Differentiating Herpes Simplex Virus Type 1 Strains. Journal of Clinical Microbiology, 47(3), 533–540. 10.1128/JCM.01565-08

Dégremont, L. (2013). Size and genotype affect resistance to mortality caused by OsHV-1 in Crassostrea gigas. Aquaculture, 416–417, 129–134. 10.1016/j.aquaculture.2013.09.011

Dellicour, S., Hong, S. L., Hill, V., Dimartino, D., Marier, C., Zappile, P., Harkins, G. W., Lemey, P., Baele, G., Duerr, R., & Heguy, A. (2023). Variant-specific introduction and dispersal dynamics of SARS-CoV-2 in New York City – from Alpha to Omicron. PLOS Pathogens, 19(4), e1011348. 10.1371/journal.ppat.1011348

Delmotte, J., Pelletier, C., Morga, B., Galinier, R., Petton, B., Lamy, J.-B., Kaltz, O., Avarre, J.-C., Jacquot, M., Montagnani, C., & Escoubas, J.-M. (2022). Genetic diversity and connectivity of the Ostreid herpesvirus 1 populations in France: A first attempt to phylogeographic inference for a marine mollusc disease. Virus Evolution, 8(1), veac039. 10.1093/ve/veac039

Dotto-Maurel, A., Pelletier, C., Degremont, L., Heurtebise, S., Arzul, I., Morga, B., & Chevignon, G. (2025). Evaluation of long-read sequencing for Ostreid herpesvirus type 1 genome characterization from *Magallana gigas* infected tissues. *Microbiology Spectrum*, e02082–24. 10.1128/spectrum.02082-24

Dotto-Maurel, A., Pelletier, C., Morga, B., Jacquot, M., Faury, N., Dégremont, L., Bereszczynki, M., Delmotte, J., Escoubas, J.-M., & Chevignon, G. (2022). Evaluation of tangential flow filtration coupled to long-read sequencing for ostreid herpesvirus type 1 genome assembly. Microbial Genomics, 8(11). 10.1099/mgen.0.000895

Drummond, A. J., & Rambaut, A. (2007). BEAST: Bayesian evolutionary analysis by sampling trees. BMC Evolutionary Biology, 7(1), 214. 10.1186/1471-2148-7-214

Farley, C. A., Banfield, W. G., Kasnic, G., & Foster, W. S. (1972). Oyster Herpes-Type Virus. Science, 178(4062), Article 4062. 10.1126/science.178.4062.759

Friedman, C. S., Estes, R. M., Stokes, N. A., Burge, C. A., Hargove, J. S., Barber, B. J., Elston, R. A., Burreson, E. M., & S.Reece, K. (2005). Herpes virus in juvenile Pacific oysters Crassostrea gigas from Tomales Bay, California, coincides with summer mortality episodes. Diseases of Aquatic Organisms, 63, 33–41. 10.3354/dao063033

Gámbaro, F., Layan, M., Baele, G., Vrancken, B., & Dellicour, S. (2025). Navigating sampling bias in discrete phylogeographic analysis: Assessing the performance of an adjusted Bayes factor.

Geay, A., & Mille, D. (2008). *ÉVALUATION PRÉCOCE DU CAPTAGE DE L’HUÎTRE CREUSE EN CHARENTE-MARITIME EN* 2008. Centre Régional d’Expérimentation et d’Application Aquacole.

Grijalva-Chon, J. M., Castro-Longoria, R., Ramos-Paredes, J., Enríquez-Espinoza, T. L., & Mendoza-Cano, F. (2013). Detection of a new OsHV-1 DNA strain in the healthy Pacific oyster, *Crassostrea gigas* Thunberg, from the Gulf of California. Journal of Fish Diseases, n/a-n/a. 10.1111/jfd.12028

Grizel, H., & Heral, M. (1991). Introduction into France of the Japanese oyster (Crassostrea gigas). Journal Du Conseil - Conseil International Pour l’Exploration de La Mer, (47), Article 47.

Houldcroft, C. J., Beale, M. A., & Breuer, J. (2017). Clinical and biological insights from viral genome sequencing. Nature Reviews Microbiology, 15(3), Article 3. 10.1038/nrmicro.2016.182

Hwang, J. Y., Park, J. J., Yu, H. J., Hur, Y. B., Arzul, I., Couraleau, Y., & Park, M. A. (2013). Ostreid herpesvirus 1 infection in farmed Pacific oyster larvae *Crassostrea gigas* (Thunberg) in Korea. Journal of Fish Diseases, n/a-n/a. 10.1111/jfd.12093

Jenkins, C., Hick, P., Gabor, M., Spiers, Z., Fell, S., Gu, X., Read, A., Go, J., Dove, M., O’Connor, W., Kirkland, P., & Frances, J. (2013). Identification and characterisation of an ostreid herpesvirus-1 microvariant (OsHV-1 µ-var) in Crassostrea gigas (Pacific oysters) in Australia. Diseases of Aquatic Organisms, 105(2), Article 2. 10.3354/dao02623

Katoh, K., Misawa, K., Kuma, K., & Miyata, T. (2002). MAFFT: A novel method for rapid multiple sequence alignment based on fast Fourier transform. Nucleic Acids Research, 30(14), Article 14. 10.1093/nar/gkf436

Keeling, M. J., Woolhouse, M. E. J., Shaw, D. J., Matthews, L., Chase-Topping, M., Haydon, D. T., Cornell, S. J., Kappey, J., Wilesmith, J., & Grenfell, B. T. (2001). Dynamics of the 2001 UK Foot and Mouth Epidemic: Stochastic Dispersal in a Heterogeneous Landscape. Science, 294(5543), 813–817. 10.1126/science.1065973

Kennedy, D. A., Kurath, G., Brito, I. L., Purcell, M. K., Read, A. F., Winton, J. R., & Wargo, A. R. (2016). Potential drivers of virulence evolution in aquaculture. Evolutionary Applications, 9(2), 344–354. 10.1111/eva.12342

Langmead, B., & Salzberg, S. L. (2012). Fast gapped-read alignment with Bowtie 2. Nature Methods, 9(4), Article 4. 10.1038/nmeth.1923

Lanier, H. C., & Knowles, L. L. (2012). Is Recombination a Problem for Species-Tree Analyses? Systematic Biology, 61(4), 691–701. 10.1093/sysbio/syr128

LaTourrette, K., & Garcia-Ruiz, H. (2022). Determinants of Virus Variation, Evolution, and Host Adaptation. Pathogens, 11(9), 1039. 10.3390/pathogens11091039

Leigh, J. W., & Bryant, D. (2015). popart: Full-feature software for haplotype network construction. Methods in Ecology and Evolution, 6(9), 1110–1116. 10.1111/2041-210X.12410

Lupo, C., Ezanno, P., Arzul, I., Garcia, C., Jadot, C., Joly, J.-P., Renault, T., & Bareille, N. (2016). How network analysis of oyster movements can improve surveillance and control programs of infectious diseases? Frontiers in Veterinary Science, 3. 10.3389/conf.FVETS.2016.02.00044

Lynch, S. A., Carlsson, J., Reilly, A. O., Cotter, E., & Culloty, S. C. (2012). A previously undescribed ostreid herpes virus 1 (OsHV-1) genotype detected in the pacific oyster, Crassostrea gigas, in Ireland. Parasitology, 139(12), Article 12. 10.1017/S0031182012000881

Martenot, C., Oden, E., Travaillé, E., Malas, J.-P., & Houssin, M. (2011). Detection of different variants of Ostreid Herpesvirus 1 in the Pacific oyster, Crassostrea gigas between 2008 and 2010. Virus Research, 160(1–2), Article 1–2. 10.1016/j.virusres.2011.04.012

Minin, V. N., & Suchard, M. A. (2008a). Counting labeled transitions in continuous-time Markov models of evolution. Journal of Mathematical Biology, 56(3), 391–412. 10.1007/s00285-007-0120-8

Minin, V. N., & Suchard, M. A. (2008b). Fast, accurate and simulation-free stochastic mapping. Philosophical Transactions of the Royal Society B: Biological Sciences, 363(1512), 3985–3995. 10.1098/rstb.2008.0176

Montoya, V., McLaughlin, A., Mordecai, G. J., Miller, R. L., & Joy, J. B. (2021). Variable routes to genomic and host adaptation among coronaviruses. Journal of Evolutionary Biology, 34(6), 924–936. 10.1111/jeb.13771

Morga, B., Jacquot, M., Pelletier, C., Chevignon, G., Dégremont, L., Biétry, A., Pepin, J.-F., Heurtebise, S., Escoubas, J.-M., Bean, T. P., Rosani, U., Bai, C.-M., Renault, T., & Lamy, J.-B. (2021). Genomic Diversity of the Ostreid Herpesvirus Type 1 Across Time and Location and Among Host Species. Frontiers in Microbiology, 12, 711377. 10.3389/fmicb.2021.711377

Morse, S. S. (2004). Factors and determinants of disease emergence. 23(2), 443. 10.20506/rst.23.2.1494

Moss, J., Burreson, E., Cordes, J., Dungan, C., Brown, G., Wang, A., Wu, X., & Reece, K. (2007). Pathogens in Crassostrea ariakensis and other Asian oyster species: Implications for non-native oyster introduction to Chesapeake Bay. Diseases of Aquatic Organisms, 77, 207–223. 10.3354/dao01829

Nater, A., Burri, R., Kawakami, T., Smeds, L., & Ellegren, H. (2015). Resolving Evolutionary Relationships in Closely Related Species with Whole-Genome Sequencing Data. Systematic Biology, 64(6), 1000–1017. 10.1093/sysbio/syv045

Nell, J. A. (2001). The History of Oyster Farming in Australia. Marine Fisheries Review.

Nicolas, J.-L., Comps, M., & Cochennec, N. (1992). Herpes-like virus infecting Pacific oyster larvae. (No. 1). 12(1), Article 1.

Norberg, P., Kasubi, M. J., Haarr, L., Bergström, T., & Liljeqvist, J.-Å. (2007). Divergence and Recombination of Clinical Herpes Simplex Virus Type 2 Isolates. Journal of Virology, 81(23), 13158–13167. 10.1128/JVI.01310-07

Ortigas-Vasquez, A., & Szpara, M. (2024). Embracing Complexity: What Novel Sequencing Methods Are Teaching Us About Herpesvirus Genomic Diversity. Annual Review of Virology, 11(1), 67–87. 10.1146/annurev-virology-100422-010336

Page, A. J., Taylor, B., Delaney, A. J., Soares, J., Seemann, T., Keane, J. A., & Harris, S. R. (2016). SNP-sites: Rapid efficient extraction of SNPs from multi-FASTA alignments. Microbial Genomics, 2(4). 10.1099/mgen.0.000056

Peeler, E. J., Allan Reese, R., Cheslett, D. L., Geoghegan, F., Power, A., & Thrush, M. A. (2012). Investigation of mortality in Pacific oysters associated with Ostreid herpesvirus-1μVar in the Republic of Ireland in 2009. Preventive Veterinary Medicine, 105(1–2), Article 1–2. 10.1016/j.prevetmed.2012.02.001

Pelletier, C., Chevignon, G., Faury, N., Arzul, I., Garcia, C., Chollet, B., Renault, T., Morga, B., & Jacquot, M. (2025). Phylogenomic evidence for host specialization and genetic divergence in OsHV-1 infecting *Magallana gigas* and *Ostrea edulis*. Infection, Genetics and Evolution, 134, 105803. 10.1016/j.meegid.2025.105803

Pepin, J. F., Riou, A., & Renault, T. (2008). Rapid and sensitive detection of ostreid herpesvirus 1 in oyster samples by real-time PCR. Journal of Virological Methods, 149(2), Article 2. 10.1016/j.jviromet.2008.01.022

Pérez-Losada, M., Arenas, M., Galán, J. C., Palero, F., & González-Candelas, F. (2015). Recombination in viruses: Mechanisms, methods of study, and evolutionary consequences. *Infection*, Genetics and Evolution, 30, 296–307. 10.1016/j.meegid.2014.12.022

Pernet, F., Lupo, C., Bacher, C., & Whittington, R. J. (2016). Infectious diseases in oyster aquaculture require a new integrated approach. Philosophical Transactions of the Royal Society B: Biological Sciences, 371(1689), 20150213. 10.1098/rstb.2015.0213

Porras-Hurtado, L., Ruiz, Y., Santos, C., Phillips, C., Carracedo, Á., & Lareu, M. (2013). An overview of STRUCTURE: Applications, parameter settings, and supporting software. Frontiers in Genetics, 4. https://www.frontiersin.org/articles/10.3389/fgene.2013.00098

Pritchard, J. K., Stephens, M., & Donnelly, P. (2000). Inference of Population Structure Using Multilocus Genotype Data. Genetics, 155(2), 945–959. 10.1093/genetics/155.2.945

Raj, A., Stephens, M., & Pritchard, J. K. (2013). Variational Inference of Population Structure in Large SNP Datasets (p. 001073). bioRxiv. 10.1101/001073

Rambaut, A., Drummond, A. J., Xie, D., Baele, G., & Suchard, M. A. (2018). Posterior Summarization in Bayesian Phylogenetics Using Tracer 1.7. Systematic Biology, 67(5), 901–904. 10.1093/sysbio/syy032

Ramchandani, M. S., Jing, L., Russell, R. M., Tran, T., Laing, K. J., Magaret, A. S., Selke, S., Cheng, A., Huang, M.-L., Xie, H., Strachan, E., Greninger, A. L., Roychoudhury, P., Jerome, K. R., Wald, A., & Koelle, D. M. (2019). Viral Genetics Modulate Orolabial Herpes Simplex Virus Type 1 Shedding in Humans. The Journal of Infectious Diseases, 219(7), 1058–1066. 10.1093/infdis/jiy631

Rannala, B. (2025). Recombination and phylogenetic inference. Evolutionary Journal of the Linnean Society, 4(1), kzaf016. 10.1093/evolinnean/kzaf016

Renault & Arzul. (2001). Herpes-like virus infections in hatchery-reared bivalve larvae in Europe: Specific viral DNA detection by PCR. Journal of Fish Diseases, 24(3), Article 3. 10.1046/j.1365-2761.2001.00282.x

Renault, T., Le Deuff, R.-M., Lipart, C., & Delsert, C. (2000). Development of a PCR procedure for the detection of a herpes-like virus infecting oysters in France. Journal of Virological Methods, 88(1), 41–50. 10.1016/S0166-0934(00)00175-0

Renault, T., Moreau, P., Faury, N., Pepin, J.-F., Segarra, A., & Webb, S. (2012). Analysis of Clinical Ostreid Herpesvirus 1 (Malacoherpesviridae) Specimens by Sequencing Amplified Fragments from Three Virus Genome Areas. Journal of Virology, 86(10), 5942–5947. 10.1128/JVI.06534-11

Robert, C. P., & Casella, G. (1999). The Gibbs Sampler. In C. P. Robert & G. Casella (Eds.), Monte Carlo Statistical Methods (pp. 285–361). Springer. 10.1007/978-1-4757-3071-5_7

Sanjuán, R. (2012). From Molecular Genetics to Phylodynamics: Evolutionary Relevance of Mutation Rates Across Viruses. PLOS Pathogens, 8(5), e1002685. 10.1371/journal.ppat.1002685

Sanjuán, R., Nebot, M. R., Chirico, N., Mansky, L. M., & Belshaw, R. (2010). Viral Mutation Rates. Journal of Virology, 84(19), 9733–9748. 10.1128/JVI.00694-10

Segarra, A., Pépin, J. F., Arzul, I., Morga, B., Faury, N., & Renault, T. (2010). Detection and description of a particular Ostreid herpesvirus 1 genotype associated with massive mortality outbreaks of Pacific oysters, Crassostrea gigas, in France in 2008. Virus Research, 153(1), Article 1. 10.1016/j.virusres.2010.07.011

Shimahara, Y., Kurita, J., Kiryu, I., Nishioka, T., Yuasa, K., Kawana, M., Kamaishi, T., & Oseko, N. (2012). Surveillance of Type 1 Ostreid Herpesvirus (OsHV-1) Variants in Japan. Fish Pathology, 47(4), Article 4. 10.3147/jsfp.47.129

Simon-Loriere, E., & Holmes, E. C. (2011). Why do RNA viruses recombine? Nature Reviews Microbiology, 9(8), Article 8. 10.1038/nrmicro2614

Srivastava, D., Ahmad, M. M., Shamim, M., Maurya, R., Srivastava, N., Pandey, P., Siddiqui, S., & Siddiqui, M. H. (2019). Chapter 12—Modulation of Gene Expression by Microsatellites in Microbes. In H. B. Singh, V. K. Gupta, & S. Jogaiah (Eds.), New and Future Developments in Microbial Biotechnology and Bioengineering (pp. 209–218). Elsevier. 10.1016/B978-0-444-63503-7.00012-7

Szpara, M. L., Gatherer, D., Ochoa, A., Greenbaum, B., Dolan, A., Bowden, R. J., Enquist, L. W., Legendre, M., & Davison, A. J. (2014). Evolution and Diversity in Human Herpes Simplex Virus Genomes. Journal of Virology, 88(2), 1209–1227. 10.1128/JVI.01987-13

Tamura, K., Stecher, G., & Kumar, S. (2021). MEGA11: Molecular Evolutionary Genetics Analysis Version 11. Molecular Biology and Evolution, 38(7), 3022–3027. 10.1093/molbev/msab120

Trancart, S., Tweedie, A., Liu, O., Paul-Pont, I., Hick, P., Houssin, M., & Whittington, R. J. (2022). Diversity and molecular epidemiology of Ostreid herpesvirus 1 in farmed Crassostrea gigas in Australia: Geographic clusters and implications for “microvariants” in global mortality events. Virus Research, 323, 198994. 10.1016/j.virusres.2022.198994

Trimpert, J., Groenke, N., Jenckel, M., He, S., Kunec, D., Szpara, M. L., Spatz, S. J., Osterrieder, N., & McMahon, D. P. (2017). A phylogenomic analysis of Marek’s disease virus reveals independent paths to virulence in Eurasia and North America. Evolutionary Applications, 10(10), 1091–1101. 10.1111/eva.12515

Vignuzzi, M., & López, C. B. (2019). Defective viral genomes are key drivers of the virus–host interaction. Nature Microbiology, 4(7), Article 7. 10.1038/s41564-019-0465-y

Wörner, S., Dragani, W. C., Echevarria, E. R., Carrasco, M., & Barón, P. J. (2019). An Estimation of the Possible Migration Path of the Pacific Oyster (Crassostrea gigas) Along the Northern Coast of Patagonia. Estuaries and Coasts, 42(3), 806–821. 10.1007/s12237-018-00492-z

Wu, X., Zhou, L., Zhao, X., & Tan, Z. (2014). The analysis of microsatellites and compound microsatellites in 56 complete genomes of *Herpesvirales*. Gene, 551(1), 103–109. 10.1016/j.gene.2014.08.054

Xia, J., Bai, C., Wang, C., Song, X., & Huang, J. (2015). Complete genome sequence of Ostreid herpesvirus-1 associated with mortalities of Scapharca broughtonii broodstocks. Virology Journal, 12(1), Article 1. 10.1186/s12985-015-0334-0

