## Supplemental Figure 1 for "Global diversity and dispersal routes of the *Ostreid herpesvirus type 1* infecting *Magallana gigas*"

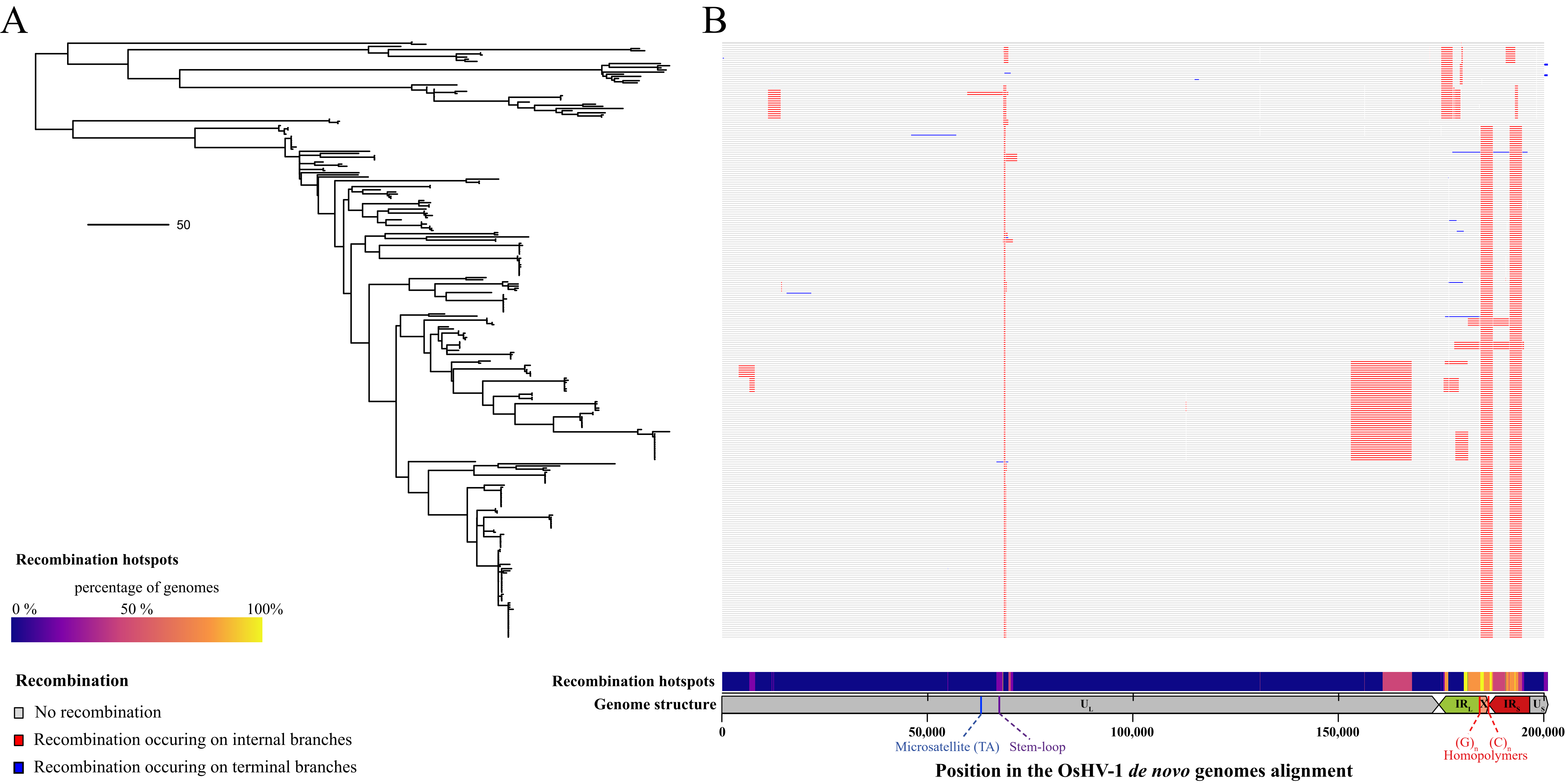


**Figure S1: Recombination events detected along OsHV-1 genomes multiple alignment.** A) Phylogenetic relationships between the 279 individuals OsHV-1 genomes estimated by a maximum likelihood approach. B) Recombination plot showing syntenic regions in grey, recombination occurring on internal branches in red and recombination occurring on terminal branches in blue for the 279 genomes. The x axis represents the position within the OsHV-1 genome. Recombination hotspots between genomes are highlighted by a gradient from dark blue to yellow.
